# Solid state NMR spectral editing of histidine, arginine and lysine using Hadamard encoding

**DOI:** 10.1101/2024.07.23.604848

**Authors:** Tata Gopinath, Alyssa Kraft, Kyungsoo Shin, Nicholas A. Wood, Francesca M. Marassi

**Affiliations:** Department of Biophysics, Medical College of Wisconsin, Milwaukee, WI 53226, USA

**Keywords:** Solid state NMR, Hadamard, sidechain, spectral aliasing, ferritin nanocage, water exchange

## Abstract

The NMR signals from protein sidechains are rich in information about intra- and inter-molecular interactions, but their detection can be complicated due to spectral overlap as well as conformational and hydrogen exchange. In this work, we demonstrate a protocol for multi-dimensional solid-state NMR spectral editing of signals from basic sidechains based on Hadamard matrix encoding. The Hadamard method acquires multi-dimensional experiments in such a way that both the backbone and under-sampled sidechain signals can be decoded for unambiguous editing in the ^15^N spectral frequency dimension. All multi-dimensional ^15^N-edited solid-state NMR experiments can be acquired using this strategy, thereby accelerating the acquisition of spectra spanning broad frequency bandwidth. Application of these methods to the ferritin nanocage, reveals signals from N atoms from His, Arg, Lys and Trp sidechains, as well as their tightly bound, ordered water molecules. The Hadamard approach adds to the arsenal of spectroscopic approaches for protein NMR signal detection.

## Introduction

The basic amino acids, His, Arg and Lys, mediate electrostatic interactions that are fundamentally important for protein structure and function. NMR spectroscopy can provide essential atomic-level structural information ^1-3^, and NMR methods have been developed to characterize the structures and dynamics of basic sidechains in solution ^4-6^ and in the solid state ^7-10^. Nevertheless, conformational exchange and water hydrogen exchange often complicate NMR studies of these basic sidechains, and challenges associated with sampling and sensitivity have limited solid-state NMR studies of basic sidechains. Since the relatively low sensitivity of ^15^N-based solid-state NMR experiments makes it time consuming to acquire separate experiments that sample either backbone or sidechain chemical shifts, it is customary to acquire ^15^N-edited multi-dimensional experiments with fast sampling that covers only the backbone ^15^N chemical shift signals (∼100-140 ppm) and excludes the sidechain ^15^N chemical shifts of His (∼160-300 ppm), Arg (∼70-95 ppm) and Lys (∼25-45 ppm). In this situation, the under-sampled sidechain frequencies are spectrally aliased upon Fourier transformation, and appear in the experimental spectral window at chemical shift frequencies that differ from their real values depending on the experimental spectral width or Nyquist frequency ^11^. Such aliased sidechain signals may overlap with other signals and complicate spectral analysis, and their phases can differ from those of backbone frequencies, resulting in mixed absorptive and dispersive line shapes.

To overcome these challenges, we present an alternative approach for solid-state NMR spectral editing of signals from basic sidechains based on Hadamard matrices ^12^. Hadamard NMR spectroscopy was originally developed to speed up multi-dimensional experiments by replacing the evolution periods of indirect dimensions with selective pulses ^13,14^, and has also been used in magnetic resonance imaging to recover aliased regions in high-resolution images ^15^. Here, we apply Hadamard encoding to deconvolute aliased peaks in multi-dimensional solid-state NMR spectra acquired at fast magic angle spinning (MAS) frequencies with ^1^H or ^13^C detection experiments.

Using the protein ferritin, we show that Hadamard spectral editing can unambiguously separate aliased and non-aliased MAS NMR signals in a single experiment. Ferritin folds as a four-helix bundle and self assembles as a highly symmetric, multimeric nanocage (**Fig. 1A**) that functions to regulate iron storage and mineralization ^16^. These properties have enabled many high-resolution structures to be determined by X-ray crystallography and cryogenic electron microscopy (EM) where it is also used as a gold standard for experimental optimization and validation ^17,18^. For these same reasons, ferritin is also ideally suited for the development and optimization of solid-state NMR experiments, where it can be readily sedimented into MAS rotors bypassing the need for crystallization ^19-21^.

**Figure 1.**
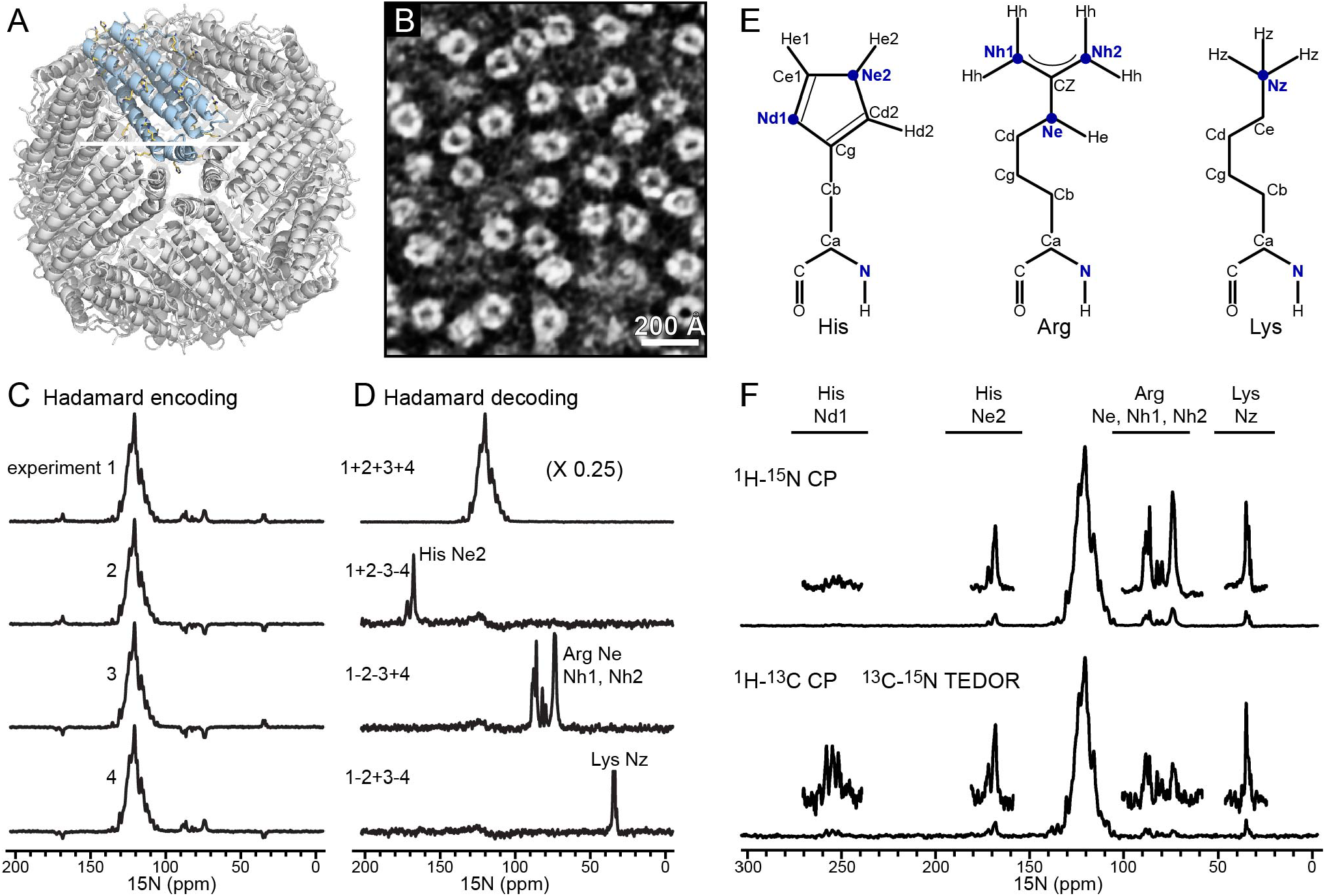
1D ^15^N Hadamard NMR spectra. **(A)** Structure of the 24-mer nanocage of mouse ferritin (PDB: 6s61) with four-helix bundle monomer in blue. **(B)** Representative transmission EM micrograph of ferritin nanocages. **(C)** 1D ^15^N spectra acquired with Hadamard encoded ^1^H-^15^N CP (**Fig. S1A**). For each Hadamard encoded experiment 1,024 transients were signal averaged. **(D)** Hadamard-decoded spectra showing spectral editing of four spectral regions. The effective number of transients for each decoded spectrum is 4,096. **(E)** Structures of the His, Arg and Lys sidechains. **(F)** ^15^N spectra obtained with ^1^H-^15^N CP (top) or _1_H-^13^C CP followed by TEDOR mixing (bottom) with 5,000 transients. Experiments performed with 60 kHz MAS.

## Results and Discussion

### Hadamard-encoded experiments

The Hadamard-encoded solid-state NMR experiments are identical to their non-Hadamard versions, except for the insertion of selective ^15^N pulses after the t1 evolution period (**Fig. S1**). The sequences include: a preparation period where polarization is transferred to ^15^N by either: ^1^H-^15^N cross polarization (CP), ^1^H-^13^C-^15^N double CP, or ^1^H-^13^C-^15^N Transferred Echo Double Resonance (TEDOR); an ^15^N or ^13^C t1 evolution period; selective 180° pulses on the ^15^N z-polarization; heteronuclear or homonuclear mixing periods; and t2 acquisition of ^1^H, ^13^C or ^15^N signal. During spectral processing, the Hadamard-encoded data are decoded for simultaneous ^15^N spectral editing of sidechain and backbone ^15^N chemical shifts (**Fig. S2**). Any ^15^N-edited solid-state NMR experiment can be acquired using this new protocol. A key advantage of the method is that the spectral encoding is obtained on ^15^N z-polarization which has T1 relaxation of the order of few seconds, thus, Hadamard spectra are acquired as efficiently as the conventional spectra without signal loss during selective pulses.

### One-dimensional spectroscopy

Each 180-residue monomer of ferritin contains ten His, eight Arg and thirteen Lys residues, and is therefore well-suited for testing basic sidechain spectral editing NMR experiments. After purification of ^15^N/^13^C labeled recombinant mouse ferritin from *E. coli*, we performed transmission EM to confirm its self-association as a symmetric nanocage of 120 Å diameter (**Fig. 1B, Fig. S3**), and then transferred the protein into the NMR rotor for MAS solid-state NMR experiments.

We first demonstrate Hadamard spectral editing with the 1D ^15^N spectra of ferritin (**Fig. 1C, D**) acquired using ^1^H-^15^N CP and ^15^N selective pulses on z-polarization followed by ^15^N acquisition (**Fig. S1A**). In addition to the high-intensity region (100-140 ppm) attributed primarily to backbone signals, three spectral regions are attributed to His Ne2, Arg Ne and Nh, and Lys Nz sidechain atoms (**Fig. 1E**). To Hadamard encode each of the spectral regions, four 1D spectra were recorded with phase encoding schemes dictated by the rows of the four-dimensional Hadamard matrix (**Fig. 1C; Fig. S2A**). In the first encoding experiment no selective pulses are applied, so the intensities of all ^15^N chemical shifts are positive. In the subsequent experiments however, a pair of selective 180° pulses are applied at the sidechain chemical shifts of Arg and Lys (experiment 2), Arg and His (experiment 3) and His and Lys (experiment 4), to invert the respective spectral regions. The phases of the His, Arg and Lys sidechain spectral regions in each scan follow the signs of the Hadamard matrix, and Hadamard transformation of the data yields the decoded spectra with unambiguous spectral editing for each sidechain type (**Fig. 1D**). Depending on the number of spectral regions sought for analysis, a suitable Hadamard matrix can be generated for ^15^N spectral editing, with the Hadamard matrix dimension defined by 2^n^, where n is a positive integer, as illustrated (**Fig. S2B**) for Hadamard editing with four- and eight-dimensional matrices ^13^.

REDOR and TEDOR recoupling sequences ^22-24^ have been successfully used to selectively filter protein signals in solid-state NMR experiments ^10,25,26^. The ^15^N sidechain spectra obtained with CP or TEDOR (**Fig. 1F**) show the utility of the latter for resolving signals from non-protonated N atoms. While the^1^H-^15^N CP signals from His Nd1 are weak or not observed at neutral pH where the site is not protonated, they are detected in the TEDOR spectrum where the initial polarization is obtained from ^1^H-^13^C CP followed by ^13^C-^15^N TEDOR mixing and ^15^N detection. In this case, the five regions of the ^15^N TEDOR spectrum can be Hadamard encoded and decoded using the eight-dimensional Hadamard matrix (**Fig. S2B**).

### Two-dimensional NH spectra

Selective encoding of ^15^N polarization in 2D NH correlation experiments (**Fig. S1B**) is achieved by applying selective 180° pulses after the ^1^H-^15^N CP and ^15^N t1 evolution periods, and before performing water suppression, reverse ^15^N-^1^H CP, and t2 ^1^H detection. Here, Hadamard encoding is implemented with four successive, interleaved scans (**Fig. S2A**) of the 2D experiment, where no selective pulses are applied in the first acquisition, and a pair of selective ^15^N 180° pulses are applied in each subsequent acquisition to invert the respective spectral regions of Arg and Lys (acquisition 2), Arg and His (acquisition 3), and His and Lys (acquisition 4). The four interleaved 2D data sets are stored in separate memory blocks and then processed with Hadamard decoding of the ^15^N dimension.

Comparison of the conventional and Hadamard NH spectra of ferritin illustrates the usefulness of the approach. The conventional NH spectrum has a number of well-resolved signals with line widths in the range of 145 Hz for ^1^H, and 49 Hz for ^15^N (**Fig. S4**) consistent with a high level of sample homogeneity ^27^. In the 40 ppm experimental spectral window of the ^15^N t1 dimension, the NH signals from basic sidechains are aliased into the amide backbone region of the spectrum, complicating their assignment to residue type (**Fig. 2A**). The Hadamard spectra, by contrast, show unambiguous spectral editing of basic sidechain signals (**Fig. 2B**). The spectra, rendered with identical intensity scale and noise floor show essentially identical peak intensities and thus indicate that Hadamard encoding does not cause signal loss.

**Figure 2.**
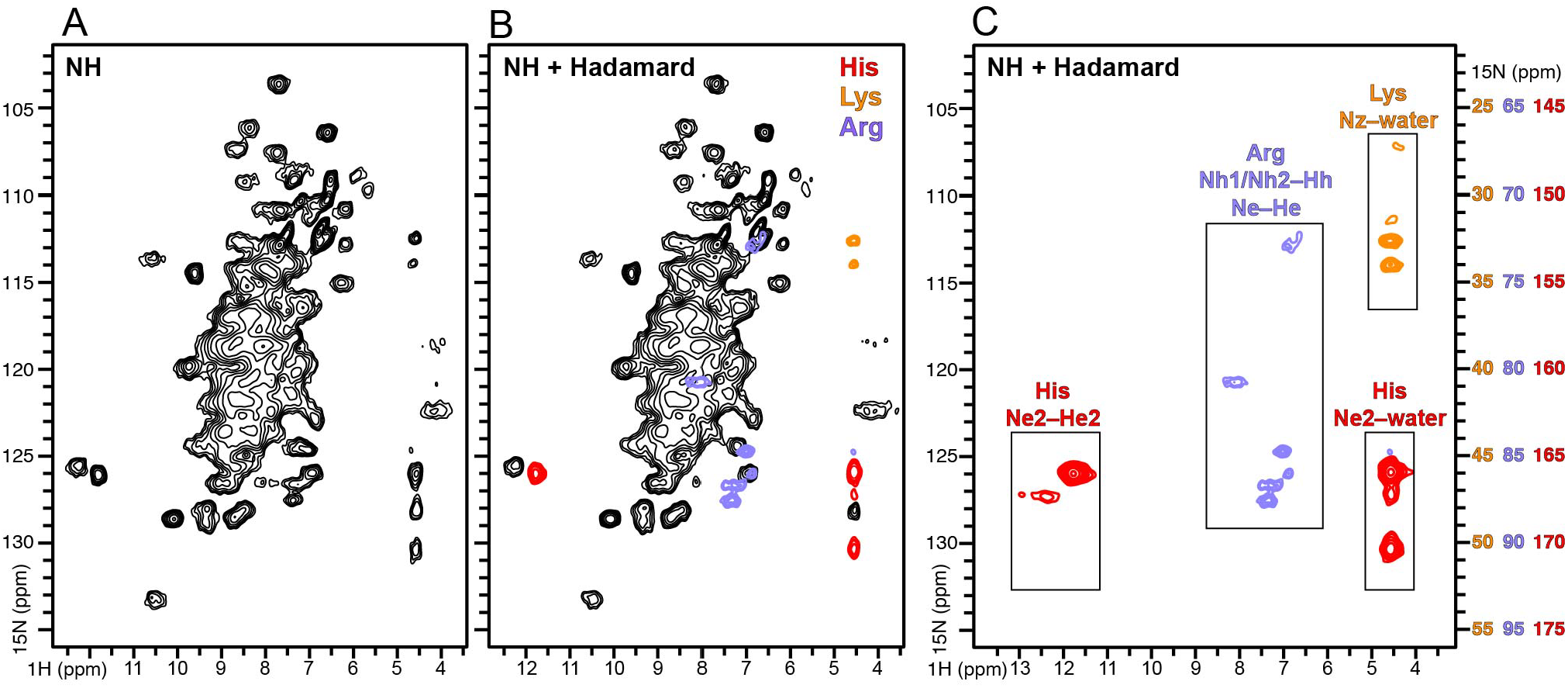
2D NH Hadamard spectra. **(A-C)** Conventional (A) and Hadamard (B, C) NH correlation spectra of ferritin, acquired at 60 kHz MAS (**Fig. S1B**). The spectra of His, Arg and Lys sidechains were separately processed and then overlayed, with ^15^N chemical shift scales corrected to their real values (colored ^15^N scales, right side). Water correlation peaks are marked for His and Lys signals. Spectra were rendered with identical scale and noise floor (A, B), or processed with window functions and phasing specific for each spectral region (C). The ^15^N chemical shift scale of His, Arg and Lys sidechains are corrected to their real values and shown in respective colors (C).

Moreover, the deconvoluted sidechain spectra can be processed with specific window functions and phase corrections optimized for each data set, a function that is not possible for convention spectral data. Thus, the Hadamard spectra processed with optimal apodization and phase corrections (see materials and methods) reveal a number of signals from His, Arg and Lys (**Fig. 2C**), including signals from Arg Nh1 and Nh2 sites which are rarely observed because hindered 180° rotations around the Ne-Cz, Cz-Nh1 and Cz-Nh2 bonds of the planar guanidinium group cause interconversions between the two Nh atoms and their attached hydrogens, in the intermediate exchange regime on the ^1^H and ^15^N chemical shift timescales ^6^.

### Detecting bound water in the ferritin nanocage

In the 2D NH spectra, signals from His Ne2 and Lys Nz sites have unique correlations to water protons (∼4.6 ppm), indicative of slow ^1^H exchange with bound water molecules on millisecond residence times ^6^. Solid-state NMR is uniquely suited for detecting bound, slow-exchange water molecules. For example, solid-state NMR revealed long-lived interactions of antimicrobial peptides with water at membrane interfaces ^28^, and demonstrated the presence of water molecules bound to key His sidechains in the influenza M2 ^1^H channel that are critical for the ^1^H conduction mechanism ^29,30^.

To confirm assignment of water correlation peaks to ferritin sidechains, we performed a 2D Hadamard NH correlation experiment with a dipolar filter where simultaneous CP from ^1^H to both ^13^C and ^15^N depletes ^1^H polarization from protein, while allowing only polarization from bound water to survive with minimal loss of signal intensity ^30^ (**Fig. S1C**). During simultaneous CP, the phase of the ^15^N radio frequency pulse is orthogonal to that of the prior CP period to prevent residual ^15^N polarization transfer back to ^1^H ^31^.

The Hadamard spectrum obtained with the water filter (**Fig. 3, blue**) has highly reduced (>90%) peak intensities compared to the non-filtered experiment, due to depletion of polarization from protein ^1^H, with a few persisting signals likely arising from ^1^H exchange with bound water. In addition, signal loss is also mediated by ^1^H T1ρ during simultaneous CP. Several signals in the Hadamard spectra of His and Lys sidechains (**Fig. 3B, 3C, blue**) resonate at the ^1^H frequency of water confirming that their corresponding N sites are water-bound and result from slow exchange with water on the millisecond time scale. In the case of Lys, the spectra indicate that the sidechain Hz protons are either in fast exchange with water and not observed, or in slow exchange and resulting in Nz-H_2_O correlation signals. Similarly for His, the strong Ne2-water correlation signals reflect bound water molecules.

**Figure 3:**
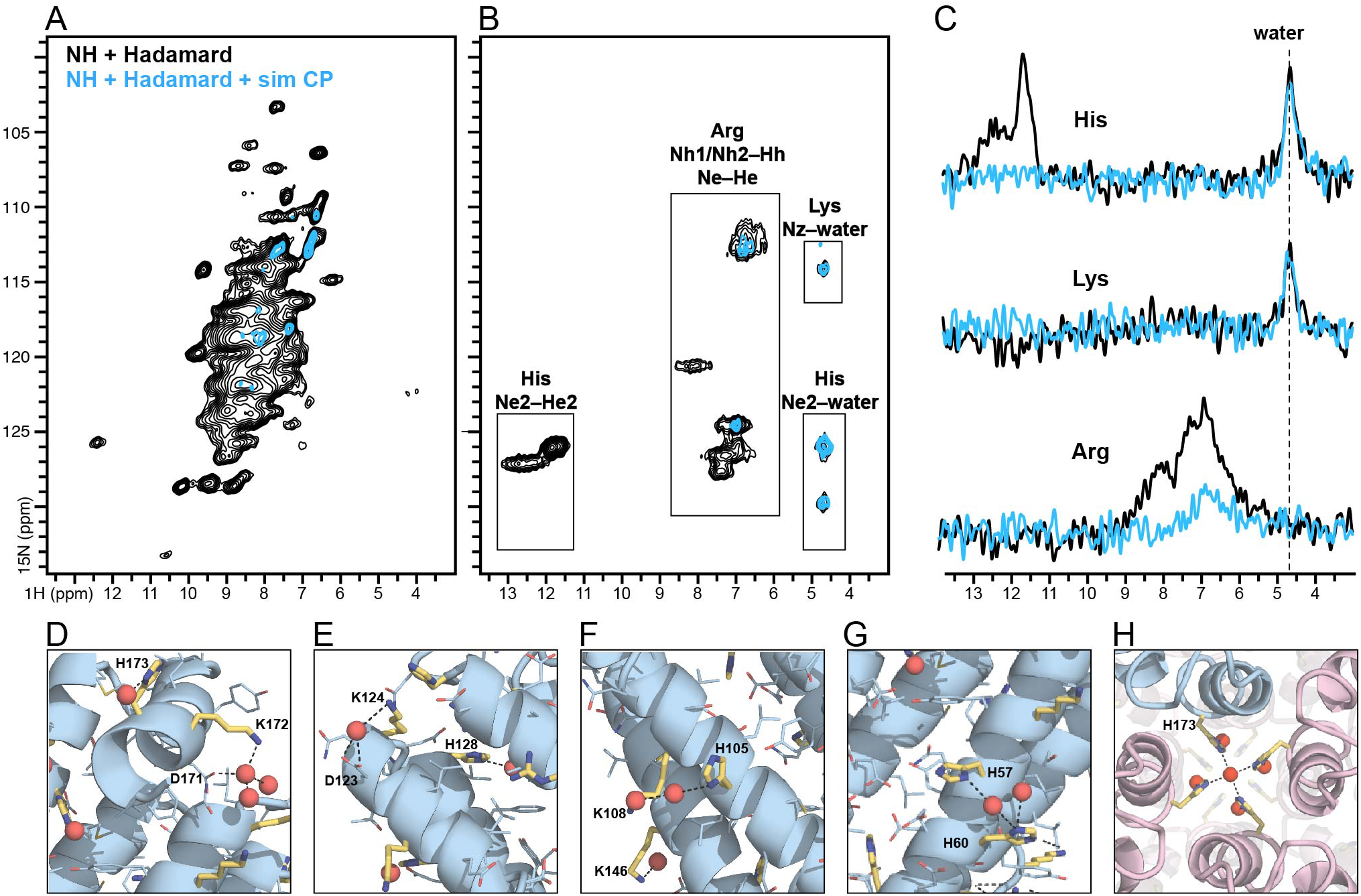
Water-filtered NH spectra. **(A-C)** Hadamard 2D NH spectra of ferritin acquired without (black) or with (blue) a simultaneous CP water filter (**Fig. S1C**) at 50 kHz MAS. Spectra were rendered with identical scale and noise floor. The spectra of His, Arg and Lys sidechains were separately processed and then overlayed (B). Projection 1D ^1^H spectra were taken from the 2D Hadamard spectra of each spectral region corresponding to His, Arg and Lys (C). **(D-H)** Examples of basic sidechains (yellow stick) bound to water oxygen (red sphere) in the structures of mouse (D-G; PDB: 6s61) or human (H; PDB: 7a6a) ferritin. Intra-molecular (D-G) and inter-molecular (H) water-mediated bridges are observed.

To examine the potential for water exchange in the context of the ferritin molecule, we analyzed the polar contacts between water O atoms and ferritin N atoms from His, Arg and Lys sidechains in the crystal structures (**Fig. 3D-H**). Several His and Lys are associated with water bridges, including K172, K124-D123 and H105-K108 on the nanocage exterior (**Fig. 3D-F**), H57-H60 on the nanocage interior (**Fig. 3G**), and the H173 tetrad which forms the three 4-fold symmetry axes of the ferritin nanocage (**Fig. 3H**). While the crystal structure does not provide information about the time scale of water exchange, our data indicate that at least some of these are sufficiently long-lived for detection by solid-state NMR. Bound, ordered water is known to play an important mechanistic role in the iron homeostatic activities of ferritin, but precisely how and where protons and water enter and exit the ferritin nanocage is an open question ^16^. The present data indicate that Hadamard solid-state NMR can offer an opportunity to contribute mechanistic information in this regard. The high quality of the ferritin NMR spectra gives confidence that resonance assignments may be obtained, paving the way for addressing such mechanistic questions.

### Two-dimensional CH correlation

TEDOR has been used as a spectral editing tool to obtain ^13^C-^15^N selective recoupling of His sidechains ^10^. Here, we combined Hadamard encoding with TEDOR to obtain ^15^N-filtered CH spectra of His, Arg and Lys sidechains (**Fig. S1E**). In this experiment, ^1^H-^13^C CP is followed by ^13^C t1 evolution, ^13^C-^15^N-^13^C TEDOR mixing, reverse ^13^C-^1^H CP, and finally ^1^H detection. Because the ^15^N magnetization is not evolved during t1, but used instead as a polarization pathway, it is possible to Hadamard encode specific C-N pairs by applying selective ^15^N pulses for specific sidechain chemical shifts. The 2D experiment was acquired with eight interleaved acquisitions and ^15^N spectral encoding corresponding to eight rows of the Hadamard matrix (**Fig. S2B**), and the eight interleaved data sets were then decoded to obtain ^15^N spectral editing of five spectral regions corresponding to the backbone and specific sidechain sites.

Comparison of conventional CH spectra (**Fig. 4A, black**), and Hadamard-encoded TEDOR-filtered CH spectra (**Fig. 4B, C, color**) enable peak assignments to specific sidechain sites. In the Arg spectrum, the weak Cz signals are most likely associated with Cz-He long range correlation since the Arg Cz atom has no directly bonded proton. In the case of His, Ne2 can be polarized by two pathways, Cd2-Ne2 and Ce1-Ne2, hence Ne2 spectral editing gives rise to two CH peaks each associated with Cd2-Hd2 and Ce1-He1 (**Fig. 4B, 4C, red**). By contrast, Nd1 can be polarized by a single, Ce1-Nd1, pathway, hence Nd1 spectral editing gives rise to a single Ce1-He1 peak for each His residue (**Fig. 4B, 4C, green**).

**Figure 4.**
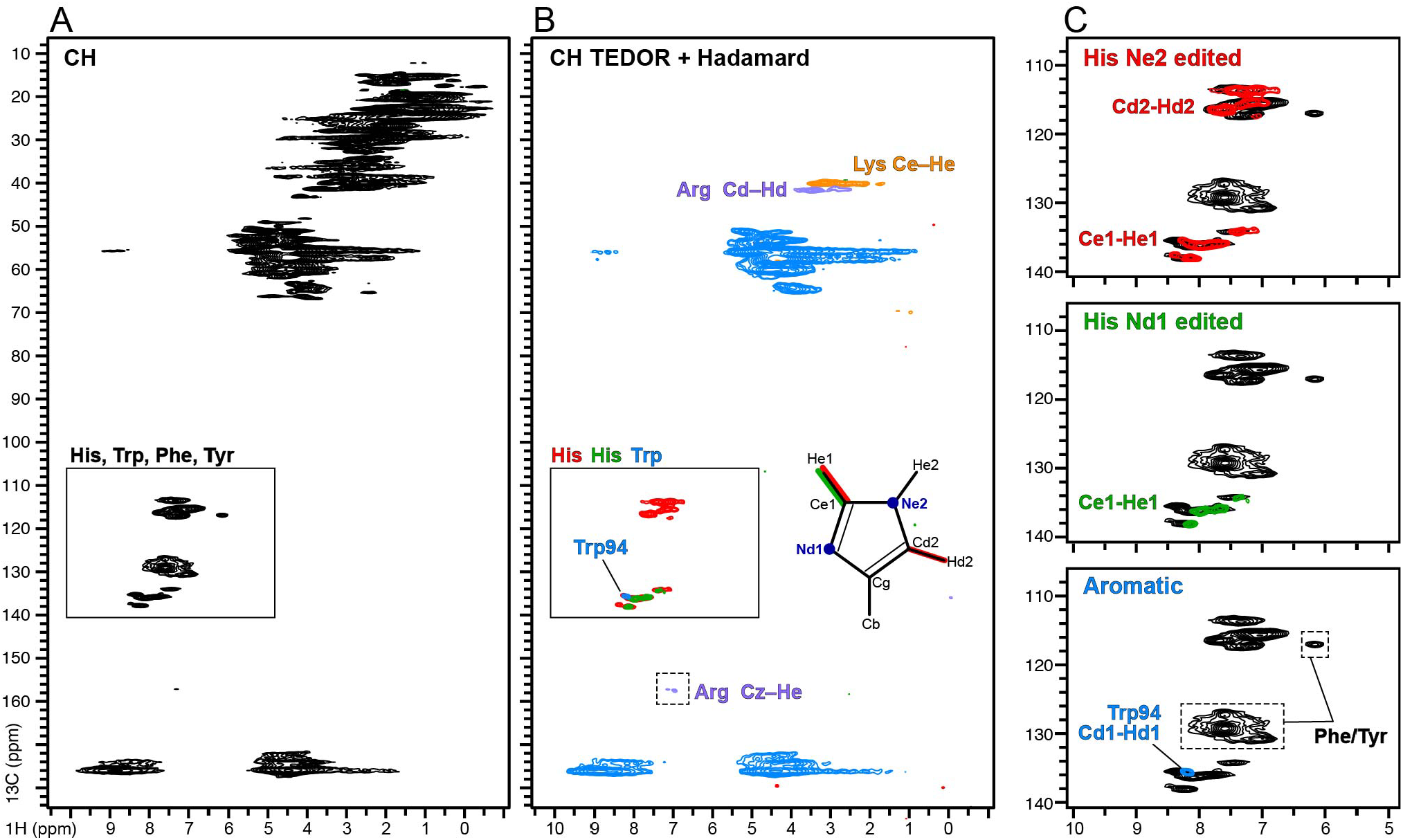
2D ^1^H-detected ^15^N-filtered CH TEDOR Hadamard spectra. **(A)** Conventional CH spectrum without Hadamard encoding. **(B)** Hadamard ^15^N-filtered TEDOR CH spectra (**Fig. S1E**). The spectra were acquired at 50 kHz MAS frequency. The Hadamard spectra of His (red, green), Arg (violet), Lys (orange) sidechains, and other N-connected C sites (blue), were processed separately and then overlayed. **(C)** Overlay of selected region of the conventional CH spectrum (A, inset box) and Hadamard TEDOR CH spectra (B, inset box).

In the aromatic ^13^C region (110-140 ppm), the TEDOR ^15^N filter recouples only His and Trp sidechain N-C pairs, whereas the N-lacking Phe and Tyr sidechains are filtered out. Thus, by separating the His sidechain signals one can unambiguously detect the sidechain Cd1-Hd1 signals from Trp94, the single Trp residue of the ferritin sequence (**Fig. 4B, 4C, blue**), while signals from the six Phe and nine Tyr can be identified by comparing the conventional non-Hadamard CH spectrum with the Hadamard ^15^N-filtered spectrum.

Finally, we implemented Hadamard spectral editing with ^13^C-detected 2D NCX and TEDOR NCX experiments (**Fig. S1D, S1F**), with spectral encoding and decoding of five spectral regions (**Fig. S2B)**. In the NCX experiment, ^15^N polarization was generated by ^1^H-^15^N CP and ^15^N correlation peaks from non-protonated N sites (His Nd1) are not observable (**Fig. 5A**). In the case of TEDOR, by contrast, the initial polarization is generated by ^1^H-^13^C CP and then transferred to ^15^N and back to ^13^C by TEDOR mixing, resulting in high intensity Nd1-Ce1 peaks (**Fig. 5B**). In total, six Nd1-Ce1 peaks and four Ne2-Ce1 peaks can be resolved for ten His in the ferritin sequence. TEDOR also yields significant intensity enhancement of all His, Arg and Lys sidechain signals compared to NCX, where lower intensity is likely due to inefficient sidechain magnetization transfer. Moreover, TEDOR offers the unique advantage of unambiguous detection of Pro signals, correlating imino N with Ca and Cd sites, while Pro signals are very weak or not detectable in NCX due to the lack of N-attached H atoms. In the case of ferritin, TEDOR enables Ca-Cd signals from the four Pro residues of ferritin to be identified (**Fig. 5B**).

**Figure 5.**
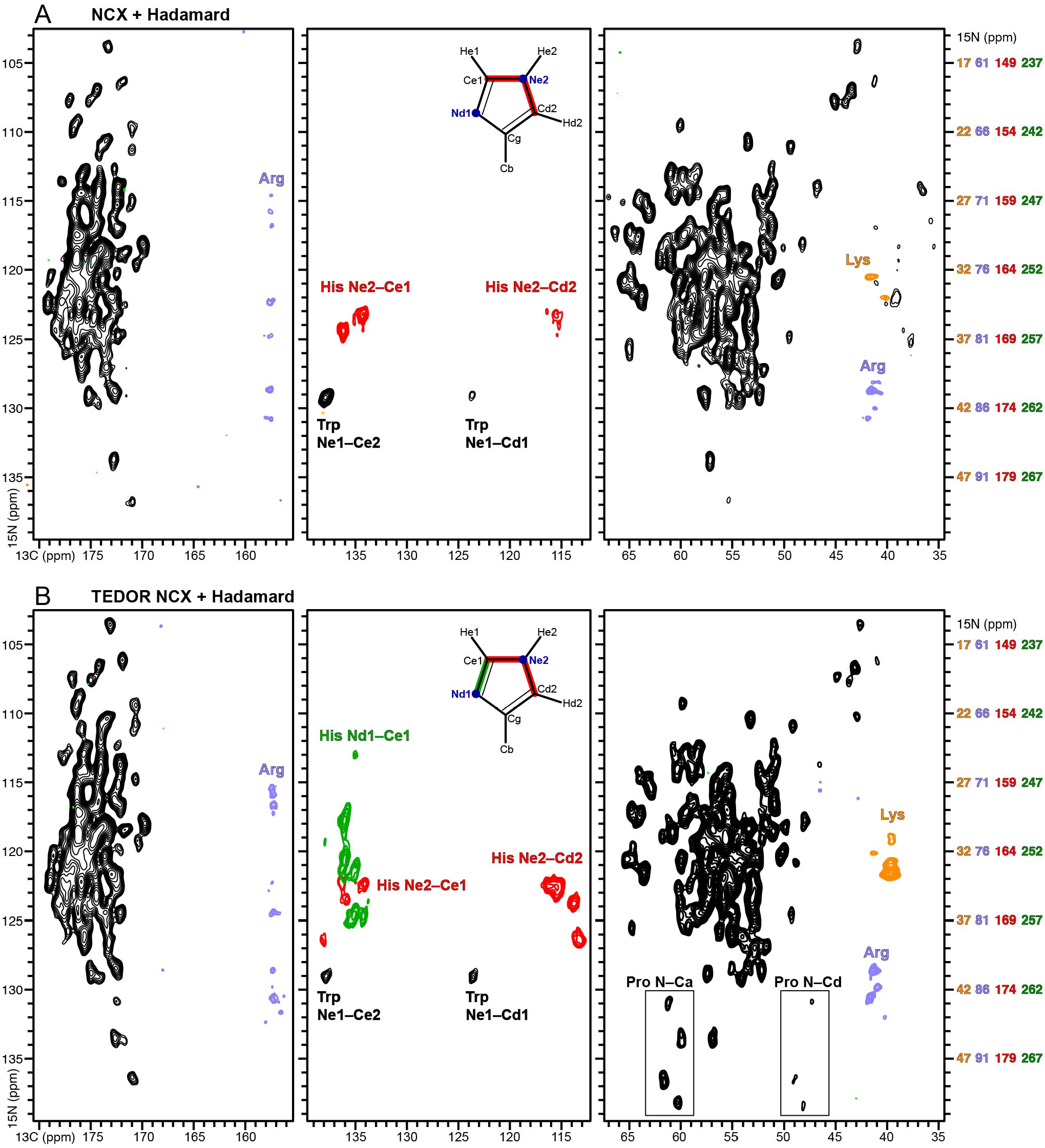
2D NCX and TEDOR NCX Hadamard spectra. The aliased sidechain spectra of His, Arg and Lys were separately processed and overlayed on the rest of the spectrum (black). The corrected chemical shift scales for sidechains are shown in respective colors. The intensity scale for His, Arg, and Lys side chain spectra was multiplied by 2 relative to the black spectrum. Spectra were acquired at 50 kHz MAS (**Fig. S1D, F**).

## Conclusions

In conclusion, we have shown that Hadamard spectral editing provides new opportunities to detect signals from backbone and basic His, Arg and Lys sidechains, which play key roles in protein structure and function. While NMR sampling of broad spectral widths increases experiment times, the Hadamard method achieves spectral editing of the entire ^15^N spectrum in a single experiment, thereby overcoming the spectral aliasing problem, without compromising spectral resolution. While demonstrated here for a series 1D and 2D experiments, Hadamard editing can be applied to a range of other 2D as well as 3D experiments. The approach is distinct from methods based on selective pulses or selective recoupling of atomic pairs, such as those used to select signals from basic sidechains, carboxyl groups of acidic residues, and aromatic residues ^7,10,26,32^. We anticipate that use of these methods with ultra-low temperature experiments, such as dynamic nuclear polarization ^33,34^, will further enhance the detection of dynamic side chains. The method, therefore, adds to the arsenal of spectroscopic approaches for protein NMR.

## Materials and Methods

### Protein preparation

Mouse apoferritin was encoded in the pET24a-mFth1 plasmid vector and transformed into BL21(DE3) *E. coli* cells (Thermo Fisher Scientific). The expression and purification protocols were adapted from Inoue and coworkers with minor modifications ^18^. Briefly, cells were grown at 37°C, in 1 L of ^13^C and ^15^N labeled M9 minimal media, up to an optical density at 600 nm (OD_600_) of 0.5, protein expression was induced by adding 1 mM IPTG, and cell culture was continued for 4 hours. Cells were sedimented by centrifugation, resuspended in buffer A (30 mM HEPES pH 7.5, 300 mM NaCl) supplemented with1 mM MgSO_4_, and then lysed with a French Press.

In a first purification step, the resulting lysate was centrifuged (20,000 g, 4°C, 30 min) to remove cellular debris, then incubated in a water bath (70°C, 10 min) and further clarified by centrifugation (20,000 g, 4°C, 30 min), and then supplemented with 330 mg/mL (NH_4_)_2_SO_4_, stirred (4°C, 10 min), and centrifuged (14,000 g, 4°C, 15 min). After dialysis (25 kDa cutoff, 4°C) against buffer A, the solution was concentrated to 500 µL (Amicon Ultra centrifugal filter; 100 kDa cutoff; Millipore-Sigma), and the protein was further purified by size exclusion chromatography in buffer A (Cytiva Superose 6 10/300 GL column).

For NMR sample preparation, the purified protein solution was first concentrated to 1 mL (Amicon Ultra centrifugal filter; 100 kDa cutoff; Millipore-Sigma), and then packed into a 1.3 mm MAS rotor (Bruker), by centrifugation (174,000 g, 4°C, 16 hr) using a spiNpack device (Giotto Biotech). This procedure yielded ∼4 mg of ^13^C/^15^N apoferritin in the rotor.

### Transmission EM

A carbon-coated copper grid (Electron Microscopy Sciences) was glow discharged using the Mini Sputter Glow Discharge System (Quorum) and 3 µL of 125 µg/mL purified ferritin solution were added to the coated side of the grid and allowed to adhere for 30 sec), before blotting residual solution with filter paper. The grid was washed and blotted twice with a water droplet before staining with a solution of 2% uranyl acetate for 30 sec. Imaging was performed on a JEM-1400FLASH (JOEL) operating at 60 kV.

Apoferritin complex sizes were estimated using the ‘analyze particles’ plugin for the Fiji platform ^35^. Images were adjusted to reduce background noise, and threshold and binary mask applied to prepare for particle selection. Particles were filtered based on circularity and minimum area to ensure accurate complex detection, and particle areas were used to back-calculate particle diameters, assuming spherical ferritin assembly.

### Solid-state NMR

Experiments were performed on a 700 MHz NeO Bruker spectrometer, equipped with a 1.3 mm ^1^H/^13^C/^15^N probe optimized for ^1^H detection. NMR spectra were processed with NMRpipe and analyzed using Poky (Sparky) software ^36,37^.

The temperature of the radio frequency (RF) coil was set to -20°C, and the effective sample temperature under spinning conditions was estimated by measuring the water ^1^H resonance frequency. Experiments were performed with spinning frequencies of 50 kHz or 60 kHz, and respective effective sample temperatures of 2°C and 18°C. During Hadamard encoding, e-BURP shaped selective pulses of 2 ms were applied at the ^15^N chemical shifts of His, Arg and Lys sidechains ^38^.

Heteronuclear decoupling was achieved using swTPPM ^39^ for ^1^H, and WALTZ-16 ^40^ for ^13^C and ^15^N, both implemented with 10 kHz RF amplitude. Acquisition times were set to 15 ms for ^1^H detection experiments, and 30 ms for ^13^C and ^15^N detection experiments. The recycle delay was set to 2 s. The 90° pulse lengths were set to 2.5 μs for ^1^H, and 5 μs for ^13^C and ^15^N. Total experiment times were: 6.2 hours for 2D NH spectra (**Fig. 2, 3**); 11.2 hours for 2D CH spectra (**Fig. 4A**); 17.8 hours for ^15^N filtered 2D CH spectra (**Fig. 4B**); and 16.3 hours for either the 2D NCX or TEDOR spectra (**Fig. 5**).

During cross polarization (CP), the ^13^C or ^15^N RF amplitudes were set to 30 kHz for experiments performed at a MAS frequency of 50 kHz, or set to 35 kHz for experiments performed at a MAS frequency of 60 kHz. The Hartmann-Hahn zero quantum conditions for ^1^H-^13^C and ^1^H-^15^N CP were optimized by varying the ^1^H RF amplitude with an 80-100% ramp ^41^, and the CP contact times set to 1 ms and 2 ms respectively.

For ^1^H-detection experiments (**Fig. 2-4**), the inverse CP, ^13^C-^1^H and ^15^N-^1^H, contact times were each set to 300 µs and 500 µs. In the water-filtered experiment, the simultaneous CP contact time was set to 500 µs. For ^13^C-detection experiments, the broadband ^15^N-^13^C matching condition was achieved with RF amplitudes of 20 kHz for ^15^N, and 35 kHz implemented with an 80-100% ramp, for ^13^C, without ^1^H decoupling ^42^. The ^13^C and ^15^N RF carriers were each set to 100 ppm and 120 ppm, and the ^15^N-^13^C contact time was set to 10 ms.

TEDOR recoupling experiments (**Fig. 4B and 5B**) were performed with 50 kHz MAS frequency, 1.44 ms (t_mix_ = 72* τ_r_) TEDOR mixing synchronized with rotor period (τ_r_ = 20 µs) with, WALTZ-16 ^1^H decoupling, and ^13^C and ^15^N 180° pulses of 6 µs ^43^. Each rotor period consists of two ^15^N 180° pulses that recouples the ^15^N-^13^C dipolar coupling. The duration of selective pulses and τ period was synchronized with rotor periods, and the z-filter Δ was set to 1 ms with ^1^H RF set to the rotor frequency of 50 kHz.

All 2D experiments were performed with States-TPPI t1 acquisition mode ^44^. The ^1^H detected experiments were performed with 200 ms water suppression, and a constant time (T-t1) period with T value set to 30 ms ^45,46^. The NH correlation experiments (**Fig. 2 and 3**) were performed by acquiring 64 transients, with 160 t1 complex points, 40.27 ppm ^15^N spectral width, and a 28 ms t1 evolution period. The CH experiment (**Fig. 4A**) was acquired with 12 transients, 1,600 t1 complex points, 284 ppm ^13^C spectral width, and a 16 ms t1 evolution period. The ^15^N-filtered CH experiment (**Fig. 4B**) was performed by acquiring 32 transients, with 1,000 t1 complex points, 284 ppm ^13^C spectral width, and a 10 ms t1 evolution period. The ^13^C-detected NCX and TEDOR experiments (**Fig. 5**) were performed by acquiring 384 transients, with 44 ppm ^15^N spectral width, 160 t1 complex points, and 25.6 ms t1 evolution time.

2D NH and CH spectra were processed using sine bell window functions (40-60 degree shifted) in both dimensions. To optimize the sensitivity of side chain spectra obtained from Hadamard decoding (**Fig. 2C, 3B**), the ^1^H dimension was processed with a Lorentz-to-Gauss window function (g1 and g2 values set to 60 and 100). The ^13^C-detected 2D spectra (**Fig. 5**) were processed with a Lorentz-to-Gauss window function (g1=60, g2=100) applied to the ^13^C dimension, and a cosine-squared bell window function applied to the ^15^N dimension. The Hadamard decoded ^13^C-detected spectra of Lys and Arg were processed with 50-100 Hz line broadening.

## Supplementary Information

Supplementary information includes Figures S1 to S4.

## Author contributions

TG and FMM wrote the main manuscript and prepared the figures. TG performed and analyzed NMR experiments. AK, NAW and KS prepared and analyzed the protein. All authors reviewed the manuscript.

## Declarations

The authors declare no competing interests.

## Statements and Declarations

The authors declare no competing interests.

