## Supporting Information for "Solid state NMR spectral editing of histidine, arginine and lysine using Hadamard encoding"

### Figures S1-S4

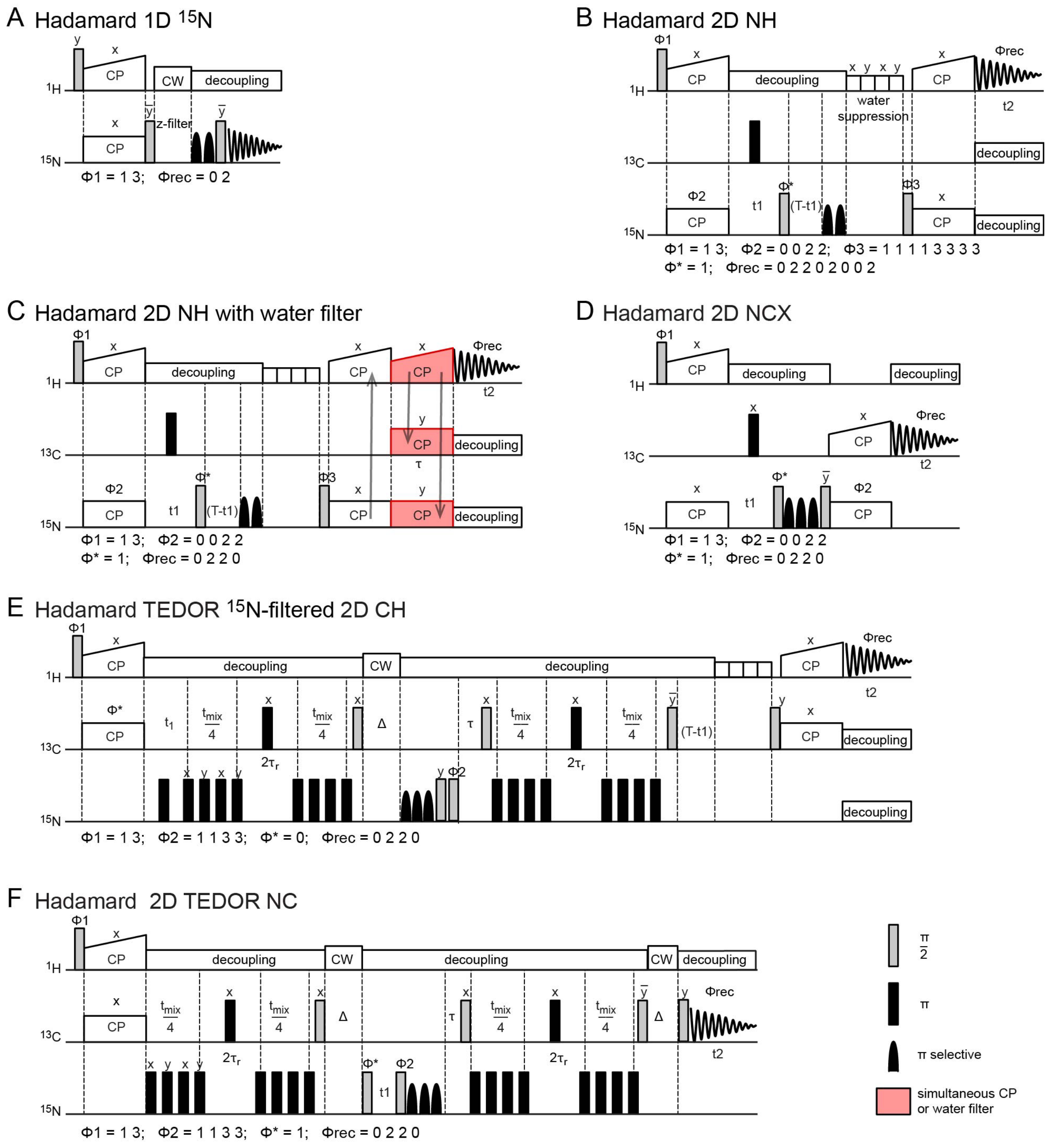


**Figure S1. Hadamard encoded 1D and 2D pulse sequences. (A)** 1D $^{15}N$. **(B)** 2D NH correlation experiment. **(C)** 2D NH with water filter. **(D)** 2D NCX. **(E)** $^{15}N$-filtered CH correlation. **(F)** 2D $^{13}C$-detected TEDOR. For 2D experiments, the t1 dimension was evolved with States-TPPI by altering the phase $\phi^*$ between x and y. In all pulse sequences Hadamard encoding of $^{15}N$ chemical shifts is obtained with four or eight interleaved scans with selective pulses applied according to the schemes prescribed by the rows of an Hadamard matrix (**Fig. S2**).

A Hadamard encoding (data acquisition)
+ (no selective pulse applied)
– (selective 180° pulse applied)

Hadamard decoding

| 15N spectral regions | His Ne2 | amide N | Arg Ne, Nh | Lys Nz |
|---|---|---|---|---|
| experiment 1 | + | + | + | + |
| 2 | + | + | – | – |
| 3 | – | + | – | + |
| 4 | – | + | + | – |

Hadamard decoding of experiments 1-4 (data acquisition)

His Ne2 → 1 + 2 – 3 – 4
amide N → 1 + 2 + 3 + 4
Arg Ne, Nh → 1 – 2 – 3 + 4
Lys Nz → 1 – 2 + 3 – 4

B Hadamard encoding (data acquisition)
+ (no selective pulse applied)
– (selective 180° pulse applied)

Hadamard decoding

| 15N spectral regions | His Nd1 | His Ne2 | amide N | Arg Ne, Nh | Lys Nz |
|---|---|---|---|---|---|
| experiment 1 | + | + | + | + | + |
| 2 | + | + | + | – | – |
| 3 | + | – | + | – | + |
| 4 | + | – | + | + | – |
| 5 | – | + | + | + | + |
| 6 | – | + | + | – | – |
| 7 | – | – | + | – | + |
| 8 | – | – | + | + | – |

Hadamard decoding of experiments 1-8 (data acquisition)

His Nd1 → 1 + 2 + 3 + 4 – 5 – 6 – 7 – 8
His Ne2 → 1 + 2 – 3 – 4 + 5 + 6 – 7 – 8
amide N → 1 + 2 + 3 + 4 + 5 + 6 + 7 + 8
Arg Ne, Nh → 1 – 2 – 3 + 4 + 5 – 6 – 7 + 8
Lys Nz → 1 – 2 + 3 – 4 + 5 – 6 + 7 – 8

**Figure S2. Procedure for Hadamard encoding and decoding.** Examples represent experiments for four (A) or five (B) $^{15}$N spectral regions using four and eight dimensional Hadamard matrices respectively. In (B) only five columns of the eight-dimensional Hadamard matrix are shown, representing five spectral regions.

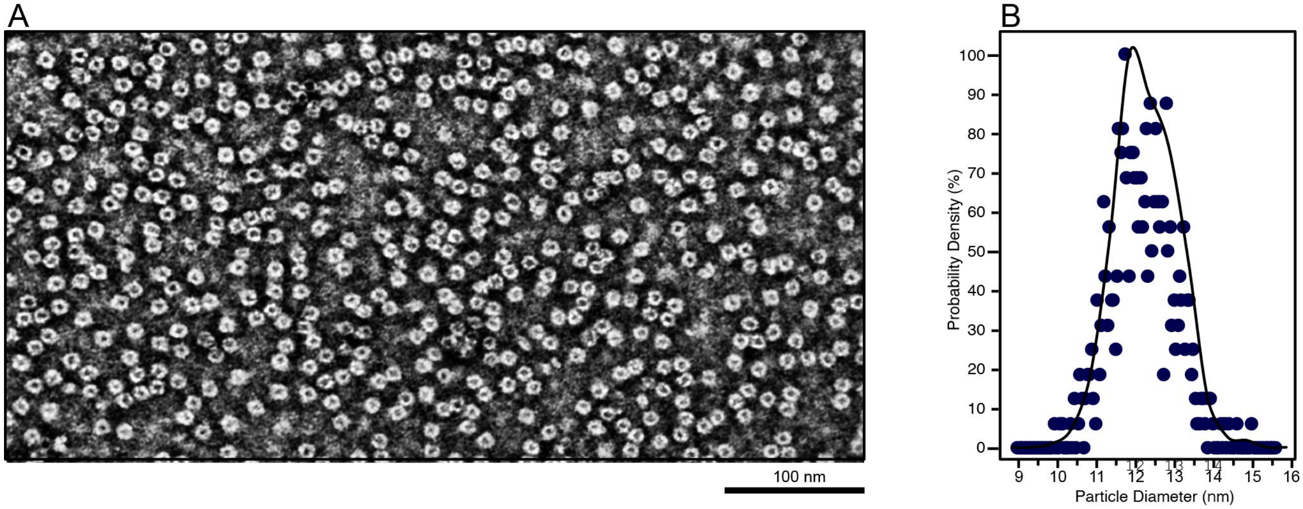


**Figure S3. TEM of recombinant mouse ferritin. (A)** Representative transmission EM micrograph of ferritin nanocages. (B) Size distribution probability density derived from particle analysis.

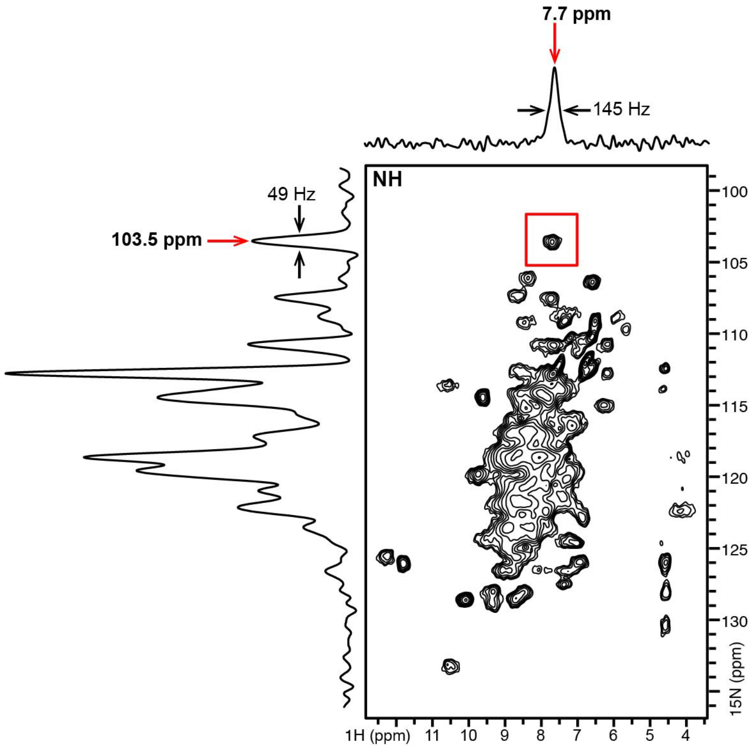


**Figure S4. 2D NH peak linewidths.** The 1D $^{1}$H and $^{15}$N spectra were taken from the 2D spectrum at 7.7 ppm and 103.5 ppm for one the Gly signals (inset red box). Individual resonances have line widths in the range of 145 Hz for $^{1}$H, and 49 Hz for $^{15}$N.
